# PeptideMind – applying machine learning algorithms to assess replicate quality in shotgun proteomic data

**DOI:** 10.1101/2020.08.20.260455

**Authors:** D.C.L. Handler, P.A. Haynes

**Author notes:** To whom correspondence should be addressed, Professor Paul A. Haynes, Department of Molecular Sciences, Macquarie University, North Ryde, NSW 2109, Australia.

## Abstract

Assessment of replicate quality is an important process for any shotgun proteomics experiment. One fundamental question in proteomics data analysis is whether any specific replicates in a set of analyses are biasing the downstream comparative quantitation. In this paper, we present an experimental method to address such a concern. PeptideMind uses a series of clustering Machine Learning algorithms to assess outliers when comparing proteomics data from two states with six replicates each. The program is a JVM native application written in the Kotlin language with Python sub-process calls to scikit-learn. By permuting the six data replicates provided into four hundred triplet non redundant pairwise comparisons, PeptideMind determines if any one replicate is biasing the downstream quantitation of the states. In addition, PeptideMind generates useful visual representations of the spread of the significance measures, allowing researchers a rapid, effective way to monitor the quality of those identified proteins found to be differentially expressed between sample states.

## Required Metadata

### Current code version

**Table 1.**
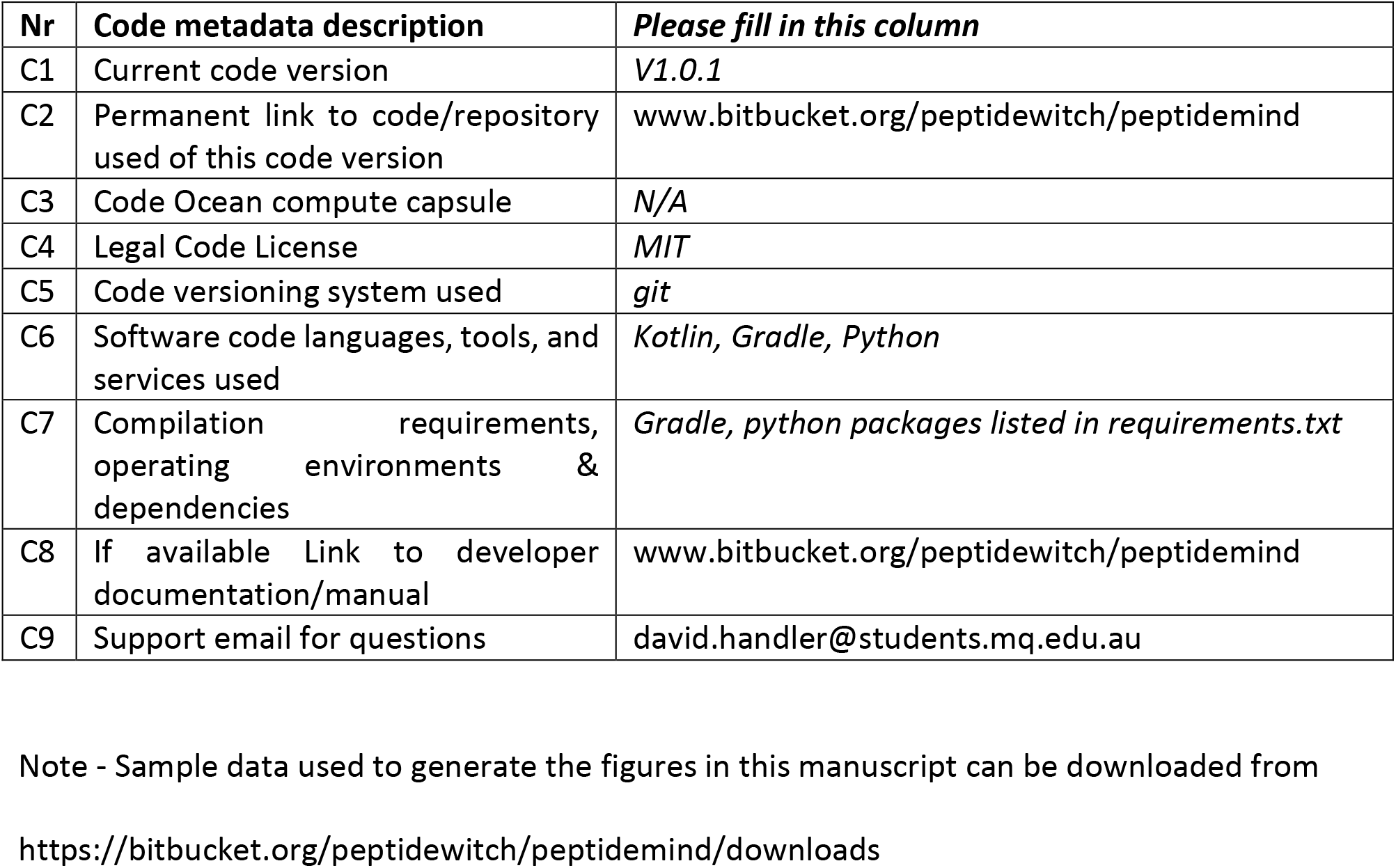
Code metadata (mandatory)

## 1. Motivation and Significance

Proteomics is the large-scale study of expressed proteins in biological systems, where the researcher undertakes analysis of biological networks that underpin cellular processes. In addition, there is a large data science component whereby raw data from a mass spectrometer has to be matched against known protein and peptide sequences before downstream analysis can occur. Making sense of all the requirements of a proteomics experiment can be a challenge; one such challenge is ensuring the validity of biological information drawn from quantitative comparisons of protein expression between test states. For example, if we have a cancer cell line dosed with a new synthetic drug, how can we effectively say that protein expression profiles generated from our treated cell state differ from those of our control cell state? There are several factors to consider, including experimental design, the process of identification of proteins, and the quantitation of proteins between these two states. Much attention has been paid to downstream forms of biological validation such as Western blotting (using an antibody to show qualitative differences between specific proteins) and parallel reaction monitoring (an orthogonal targetted mass spectrometry based method) as means to confirm an increase or decrease in protein expression between states [1]. However, there is also much to be gained from applying a more rigorous set of statistical or analytical tools prior to laboratory follow-up experiments. As Kall and colleagues describe in their seminal paper on the Percolator software [2], the application of smart methods to discriminate good quality from poor quality data can greatly improve the quality of the dataset, which can clarify any downstream quantitation performed using the data in question.

Over the past two decades, researchers have leveraged the power of machine learning algorithms (MLAs) in order to assist with shotgun proteomics data analysis. Classification algorithms are a natural fit for determining biomarker identity from state comparison experiments [3], while support vector machines have been utilised in a semi-trained fashion to help determine false discoveries in peptide to spectrum matching [2]. Machine learning algorithms have also been used to aid with peptide to spectrum matching, for example in the application of decision trees to utilise ion fragmentation patterns for peptide identification [4]. Recently, with the advent of easier implementation of MLAs via python packages such as Python’s scikit-learn, more proteomics researchers are choosing to enhance their data analysis with MLAs [5].

When it comes to their application, MLAs are by no means a silver bullet that allows a researcher to exclusively target critical insights. Not every problem can be solved by running data through Support Vector Machines or Generalised Linear Models, and not every dataset is suited to, for example, Random Forest Classification. Sometimes a simple Decision Tree can suffice. There is also the consideration of training sample sizes – the oft-quoted adage of ‘the more data, the better’ makes no *a priori* assertion regarding the quality of the data, and it may well be the case that smaller, more accurate data pools are better to train against [6]. Shotgun proteomics datasets, then, are an interesting dataset to work with, given their relatively small size compared with transcriptomics and genomics, as well as their proportion of valuable biological information relative to their noise level, or junk data. Unlike binary sentiment/natural language analysis (like understanding if a movie review is positive or negative by word choice alone), or qualitative categorisation (such as movie ratings), proteomics data is a mixture of noise and overlapping signal. A deep learning approach to six replicates of two states may not provide a clear insight into the relationship between these two states, whereas a more simple classifier-based approach could potentially offer insights as to the quality of the comparisons being made.

In this report, we introduce PeptideMind, a Kotlin/Python hybrid program that utilises a cohort of classifier MLAs to analyse replicate quality from proteomic peptide to spectrum matching search engine outputs. Following on from the same-same architecture that our lab group has developed [7], PeptideMind will take six replicates of control and treatment protein ID data and conduct multiplexed non-redundant analysis on the protein identifications (IDs). The output of the PeptideMind process is a series of graphics by which the researcher can assess overall replicate quality and check for outliers, as well as focus on specific protein identifications of interest and analyse the spread of difference in expression levels, thus assisting the researcher in providing a confident measure of statistical validation.

## 2. Software Description

PeptideMind has two components. The first is a Kotlin/Java environment that can be replicated through use of the Gradle file included within the repository. Working with a development environment such as IntelliJ by Jetbrain should handle the installation automatically; manual installation can be achieved with other environments. Regardless, Gradle is mandatory. Secondly, PeptideMind requires a version of python to be installed on the user system. Currently, PeptideMind defaults to a specific version of Python installed on the Path (a future update will allow users to point to a virtualenv such as pipenv). This python environment should have the packages installed from the requirements.txt file in the PeptideMind source directory.

The software is comprised of a single GUI page made with TornadoFX, as shown in Figure 1. Users follow the control flow from left to right hand side of the page, selecting the following elements:

1. The folder location for the control state data
2. The folder location for the treatment state data
3. The type of peptide to spectrum matching search engine used
4. The type of MLAs to use for analysis (at least 1 must be selected)
5. The scope by which the analysis is conducted on the whole dataset
6. Whether any proteins of interest should be targetted. If users input a specific text identifier (in the same format as their PSM engine from step 3) the resulting analysis will only focus on these identifiers to the exclusion of all others.
7. A folder location for the output data, and
8. A ‘start’ button

**Figure 1:**
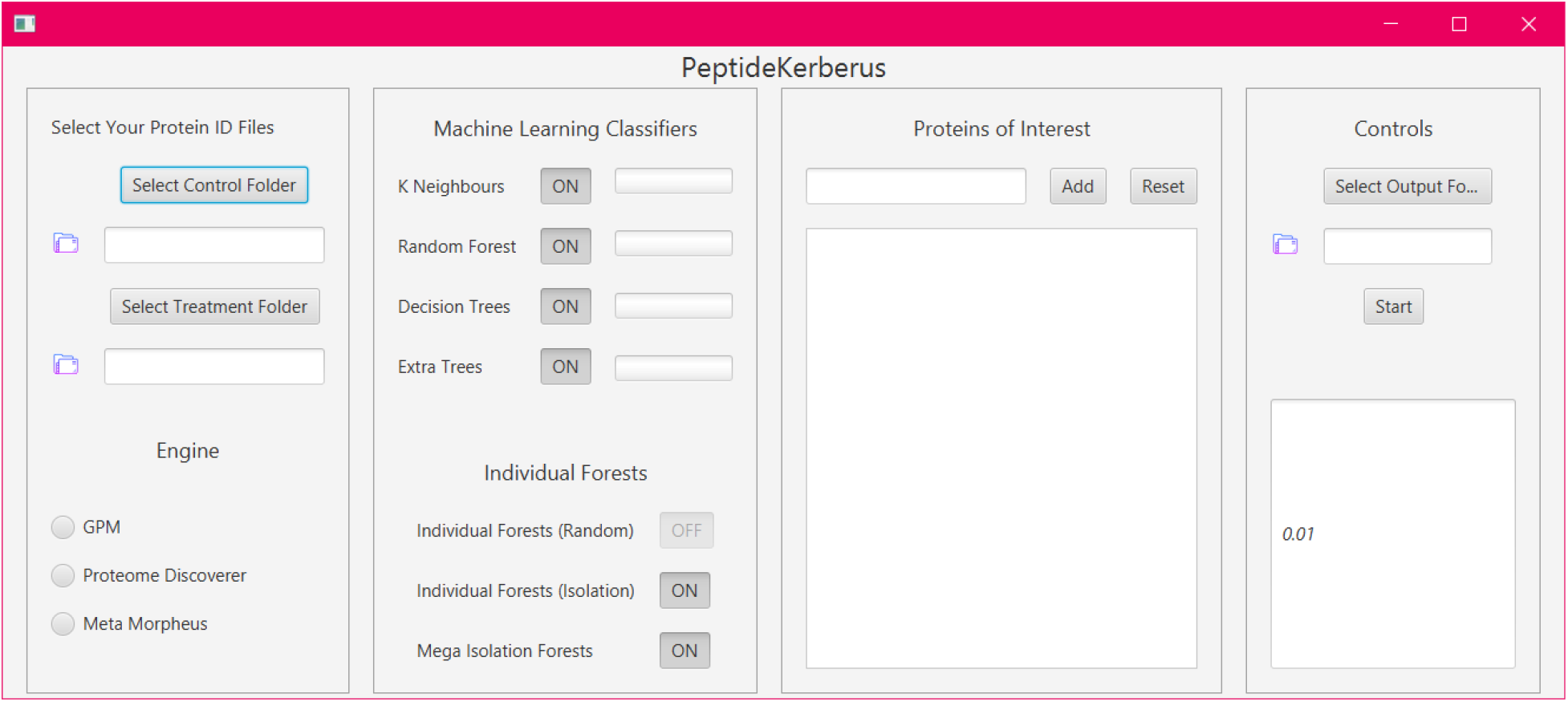
Graphical User Interface which allows interaction with the PeptideMind software.

The resulting output will be contained within the folder specified in Step 7. Users must separate their control and treatment states data into separate folders and name their replicates according to the following structure: *%name-R%replicate_number.* For example, statin-R1, statin-R2, etc, for the control state. Both states must have exactly six replicates each.

PeptideMind outputs four broad categories of results.

1. A series of excel files containing the common protein identifiers found across all six replicates for both states, including their Student’s T-Test significance values at both the spectral counting level and the exponential logarithmic normalised spectral abundance factor (NSAF) level [8,9].
2. Isolation Forests for each common ID displaying the aforementioned significance measures along the X and Y axis.
3. A MultiLabel deviation plot, where each replicate is shown against a middle line value of 0.5. This plot is designed to give users, at a glance, some idea of which replicates in which state are contributing to up- or down-regulation of significant proteins.
4. A ‘Mega Isolation’ Forest comprising an aggregate spread of protein significance measures along all four hundred combinations.

A more in-depth explanation of the processes of PeptideMind and how it arrives at the end results is provided below.

### Result category 1: .csv outputs

PeptideMind begins with two sets of sextuplicate results from proteomics peptide to spectrum matching search engines, currently including Proteome Discoverer [10], Meta Morpheus [11] and X!Tandem [12]. Broadly speaking, what the PeptideMind program aims to do is to produce an internal measure of inter-replicate variability, and display the spread of protein expression level variance, for the user to determine if subsequent differential analysis is worthwhile. To achieve this aim, the program begins by sorting the six replicates of the control and six replicates of the treatment state into sub-experiments for analysis, as shown in Figure 2. A pair of states with six replicates apiece can be split into two sets of three in four hundred non-redundant pairs. These pairs of three by three comparisons each constitute their own analysis. This pairwise combination undergoes a round of data filtering by Minimum Spectral Counting [13,14] before being subjected to two different types of Student T-Tests – one based on the spectral count of each protein, and the other based on natural log NSAF values. Any protein identifier that is considered significantly differentially expressed is recorded. This analysis is then repeated by shuffling new combinations of paired triplets between state one and state. Next, a list of proteins common to all four hundred triplet comparisons is produced, and the significance value from the two types of T-Tests are matched to the protein identifier. We then consider this data our training set for the machine learning algorithms. Data from this process is stored in .csv outputs for the user to examine for their own interest.

**Figure 2:**
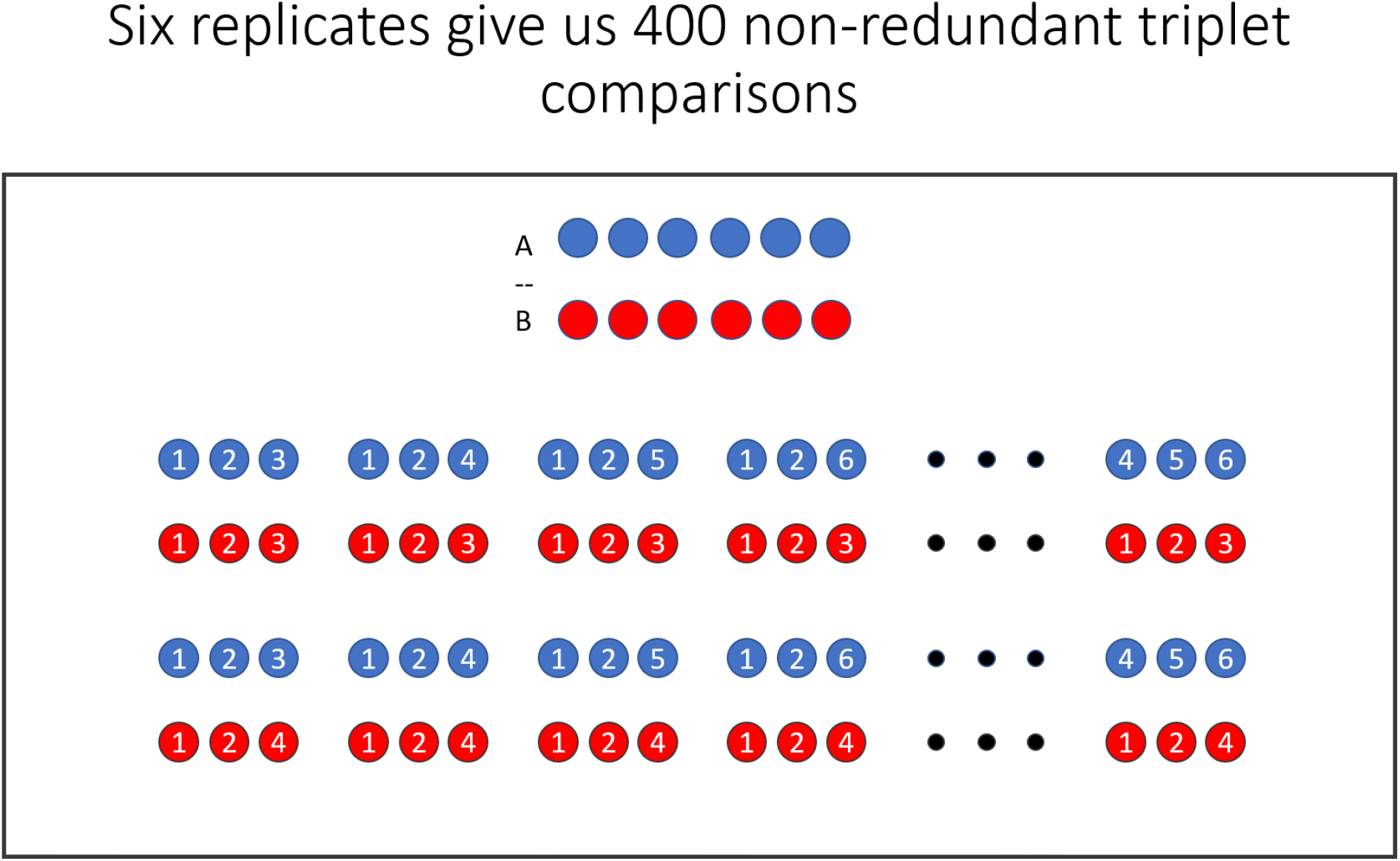
schematic diagram illustrating the replica permutation processing employed by PeptideMind.

### Result category 2: isolation forests for common protein identifiers between the states

Next, each of the common IDs from the four hundred tests is subjected to an individual Isolation Forest algorithm. The results from students T-Tests on lnNSAF and spectral count data are plotted in two dimensions, with the background color-gradient coded to correspond to the spread of significance results within the data as determined by an Isolation Forest algorithm. The results from all four hundred test combinations are shown as black dots, while a single yellow dot corresponds to the significance measure of the protein from the original 6 by 6 replicate test conducted. This serves as an anchor for the results: most researchers would only see this single measure of significance, but now, graphically, we have a way of determining the spread of significance measures in a manner that is intuitive and visually informative.

### Result category 3: multi-MLA analysis of replicate contribution to significance measures

Another useful measure of inter-replicate variability comes from a blind assignment of protein identifiers with regards to their corresponding differentiation levels from replicate values as determined by four separate multi-classifier algorithms selected by the user in Step 4 of the workflow described earlier. The results are displayed as a histogram plot which indicates the relative contribution of each replicate towards the quantitative differences of all protein identifiers shared between the states. Each data point is the average value found from the selected MLAs. What the machine learning consensus network here achieves is a blind assignment of protein identifiers to replicates, thus allowing the researcher to see if any one replicate in particular is consistently contributing more weight to differential analysis.

## 3. Illustrative Examples

To provide illustrative examples, we analysed data from two ongoing studies in our laboratory. The first study involved proteomic analysis of *Eucalytpus grandis* leaf tissue, with six replicates corresponding to young healthy leaf tissue set and six replicates from old senescent leaf tissue. Proteomic data was acquired using the X!Tandem algorithm for peptide to spectrum matching. The second study involved proteomic analysis of two different laboratory yeast strains designated CCC and CCB, with proteomic data acquired using the Meta Morpheus algorithm for peptide to spectrum matching.

The individual Isolation Forests for data generated by the program for each shared protein identifier are shown in Figure 3 for two selected proteins from the Eucalyptus experiment: K1C9 is a human keratin protein present at variable levels as a result of sample handling contamination; XP_010027978.1 is a serine hydroxymethyltransferase metabolic protein. In Figure 3a the protein ID significance shows a wide spread of results for the K1C9 protein across the different comparisons, while Figure 3b shows a very tightly correlated spread of protein ID significance for the metabolic protein. This can be viewed as a form of statistical validation; if a protein ID of interest displays a spread of significance similar to Figure 3b, rather than Figure 3a, then we can say with confidence that this result is more likely to be significant and less influenced by inter-replicate noise.

**Figure 3:**
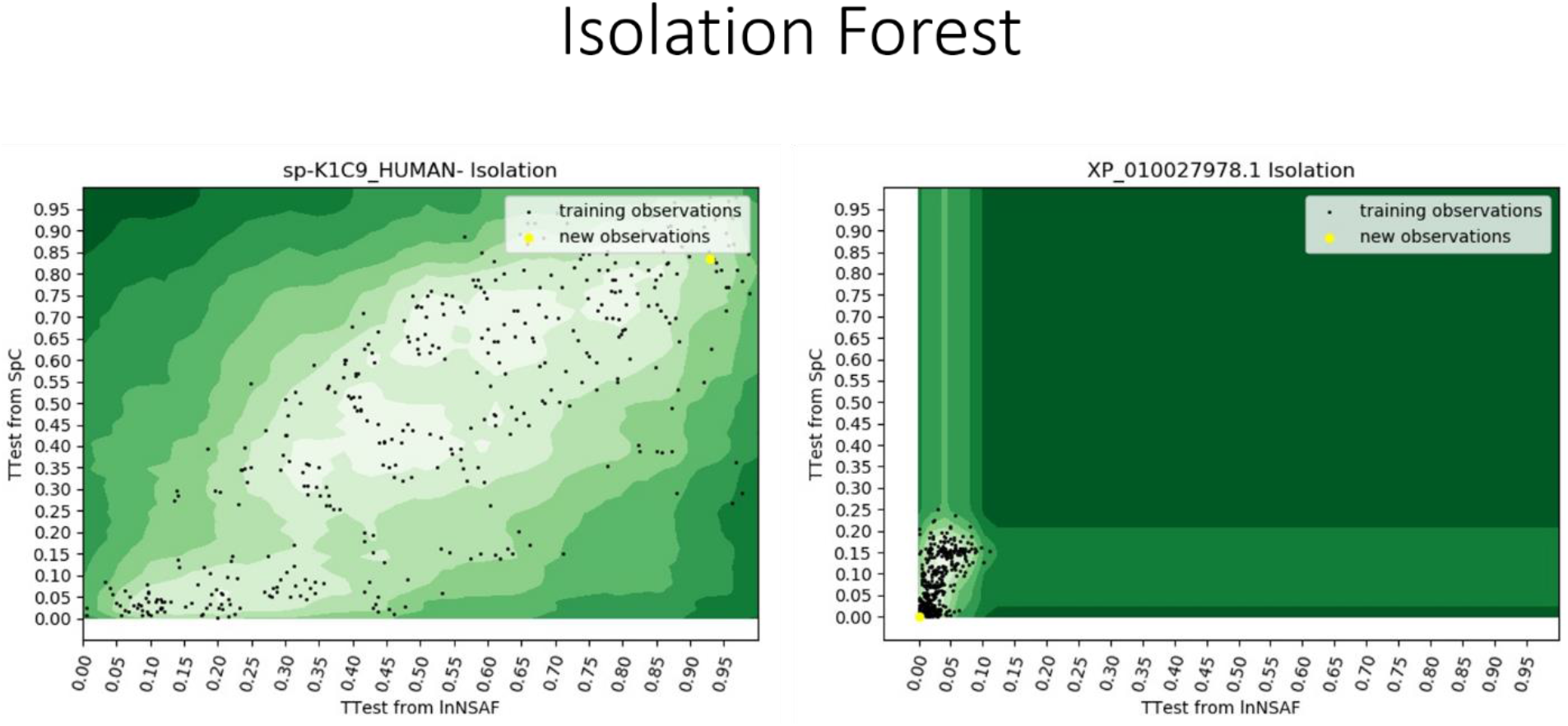
Isolation Forests generated by PeptideMind for two selected proteins from an experiment comparing the proteome of young and old eucalyptus leaves. (A) K1C9, a human keratin protein present at variable levels, (B) XP_010027978.1, a serine hydroxymethyltransferase metabolic protein.

Figure 4 shows the histogram plots of the average output values found from the selected MLAs, for both the Eucalyptus data set and the yeast data set. In Figure 4a, for the Eucalyptus data, there is clearly significant variation between the replicates. In the data from young leaf tissue (replicates R1-R6), replicates 1 and 4 contribute relatively less to the differentially regulated protein identifiers, whereas replicates 2 and 6 are overrepresented. In the data from old leaf tissue (replicates R7 to R12), replicate 8 contributes more weight to protein differentiation than replicates 10 and 12. Replicate 4, in particular, may not be truly representative of the proteome state given the relative disparity observed. In contrast, the histogram shown in Figure 4b for the yeast data indicates that all 12 replicates are internally consistent, and all contributed relatively equally to the observed differences in protein expression.

**Figure 4:**
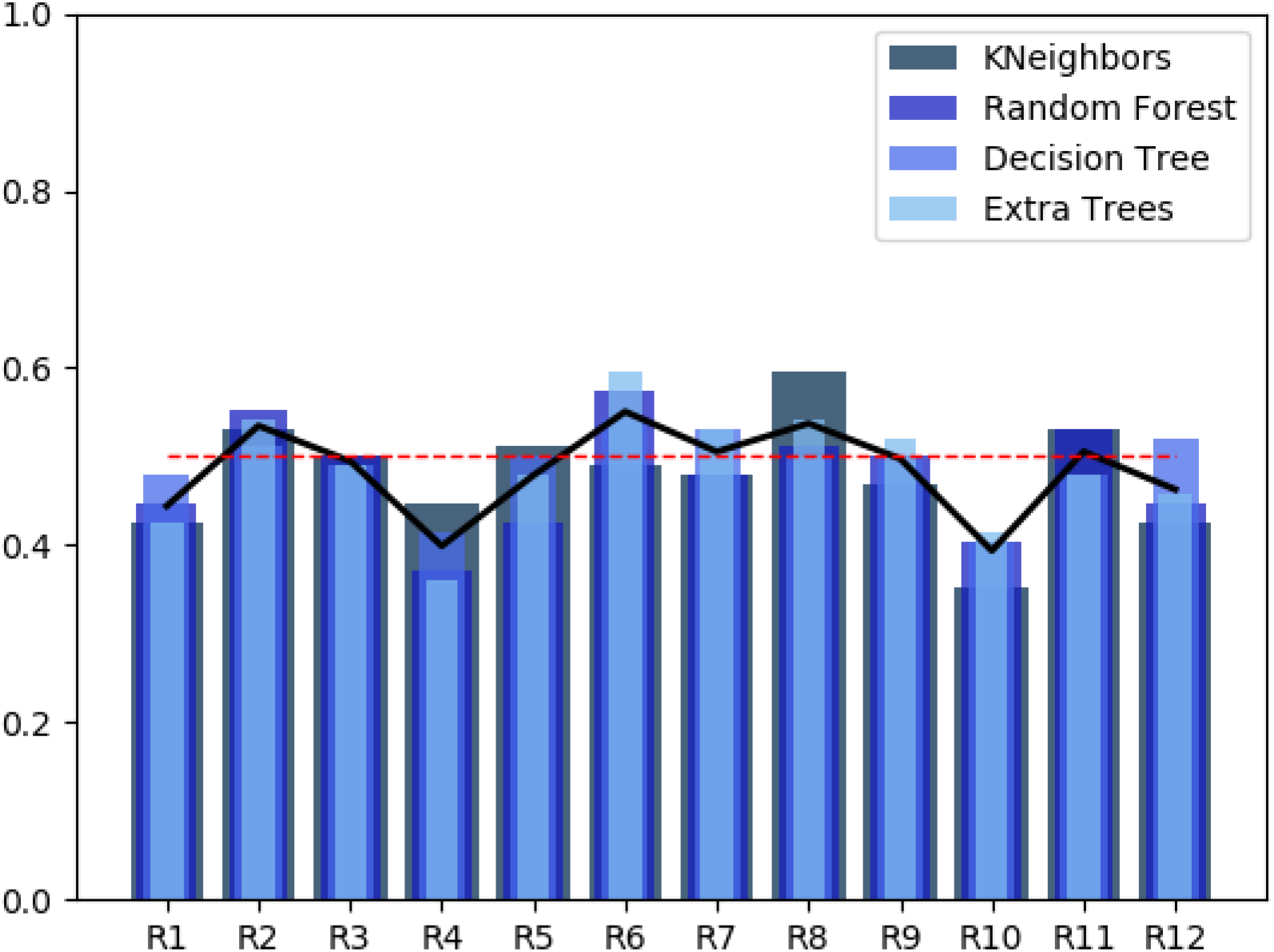
Histograms of the average output values found from the MLAs used by PeptideMind. (A) data from an experiment comparing the proteome of young (replicates R1-R6) and old (replicates R7-R12) eucalyptus leaves, (B) data from an experiment comparing the proteome of two laboratory yeast strains known as CCB (replicates R1-R6) and CCC (replicates R7-R12).

**Figure 4a.**
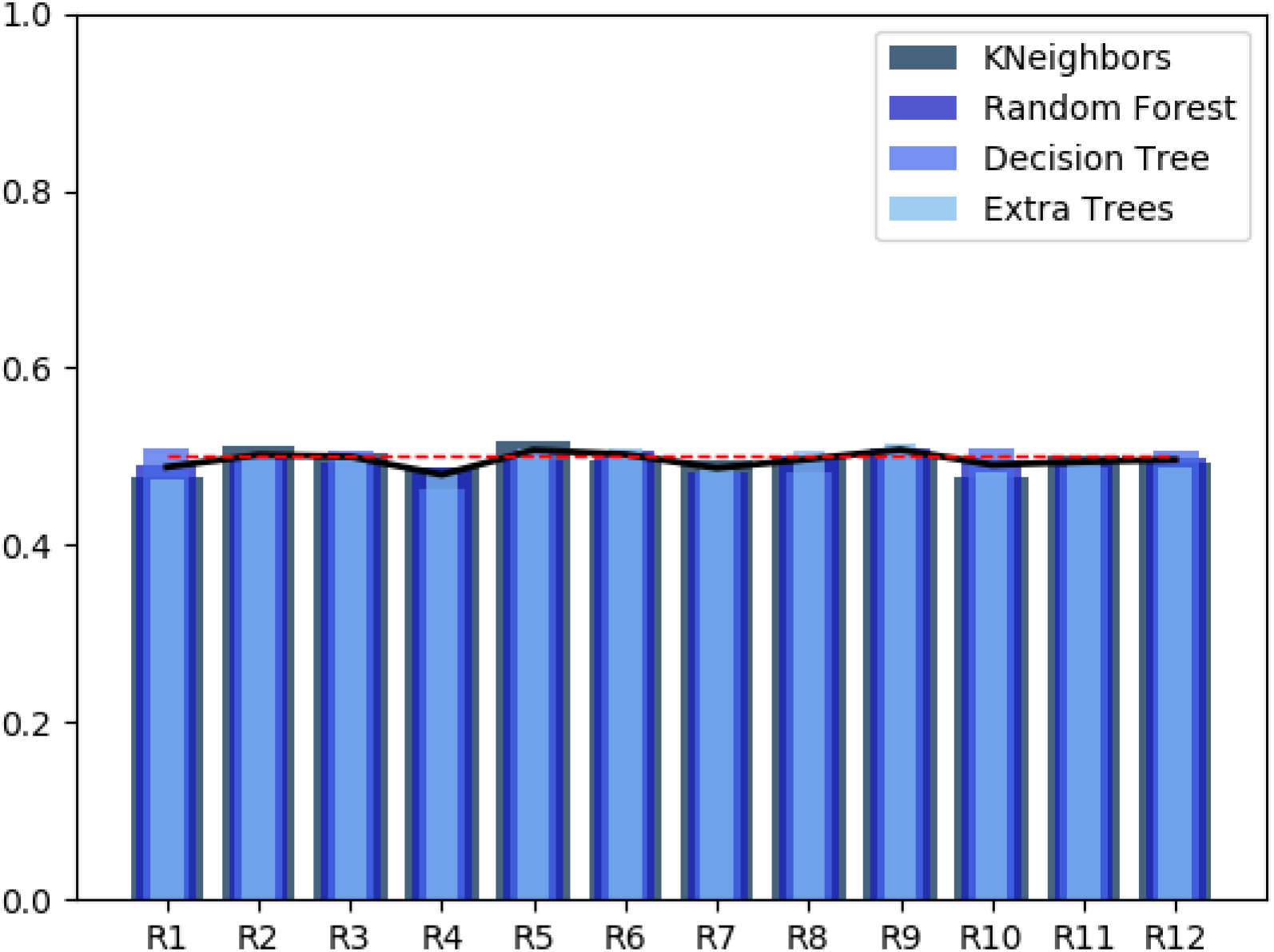

The red dotted line in the histograms represents an ideal result – if we are to assume that all of the replicates hold equal analysis weight in the course of the experiment, then every replicate should fall on the dotted line and report the same result for every one of the 400 triplet combinations. In reality this is not the case, as some replicates have higher numbers of specific proteins relative to their state cohort. This chart is a visual indicator of how biased our results are, in terms of which replicates are over- or under-represented in their contribution to protein significance spreads. A more even histogram indicates high quality replicates and low levels of variable noise, whereas a histogram more reminiscent of a city skyline may indicate significant problems in the reproducibility of the data.

## 4. Impact

PeptideMind provides user with clear visual metrics concerning the validity of their downstream quantitation profiles, by highlighting the spread of variance for every protein identifier and every replicate within the total system. At a glance, the proteomicist can understand which, if any, of their replicates are biasing the downstream results. As such, we consider PeptideMind to be a useful first step in the process of data validation prior to subsequent experiments. Consider the XP_010027979.1 metabolic protein from Figure 3b. If an additional analysis or experiment determined this protein was biologically significant within our system, we would also be able to point to this Isolation Forest result as an indicator of statistical significance that falls within expected parameters. Conversely, we would consider the K1C9 protein in Figure 3a as showing too much variance for realistic application of statistical measures of significance. Biological conclusions that rely on such a protein would be considered putative until further orthogonal validation experiments were performed.

At present, PeptideMind is not intended as a replacement for orthogonal protein validation experiments such as Western blotting, Parallel reaction monitoring, or other orthogonal mass spectrometry experiments. Rather, PeptideMind should serve as another tool in the proteomics toolbox which can be used to provide extra rigour for results, and demonstrate the validity of underlying quantitation without the need for additional experiments. The biggest hurdle with the design of this program, however, is in the requirement for two states to have six replicates each. As replicate costs can be burdensome, PeptideMind is recommended for experiments in the discovery phase where tissue can be sourced relatively cheaply. In so doing, the researcher can accumulate a solid set of data backed up by PeptideMind, and the judicial use of other statistical measures, to narrow down their list of protein identifiers for further analysis.

PeptideMind is still in development phase, and there is much room for improvement. Some future additions may include:

1. Extra permutation potential when 7, 8, 9 or more replicates are specified per state
2. Additional MLAs incorporated into the replicate bias analysis
3. The incorporation of a python virtual environment for cleaner code production
4. Better real-time feedback to the user to update what stage the program has reached.

We hope that PeptideMind may serve as the inspiration for future experimentation and software development that leverages the power of permutations with MLAs for better, more specifically tailored analysis of differentially expressed proteins in proteomics experiments.

## 5. Conclusions

In this report, we have demonstrated the capabilities of the PeptideMind software in providing a valuable tool of statistical validation for data analysis pipelines in shotgun proteomics experiments. Leveraging the power of MLAs with permutation analysis, PeptideMind is capable of generating simple yet powerful graphical metrics whereby the user can assess the quality of their replicates, differential expression profiles, and resulting quantitation. In the future, we hope to expand the capability of the platform to incorporate further improvements and additional features.

## Conflict of Interest

We confirm that there are no known conflicts of interest associated with this publication and there has been no significant financial support for this work that could have influenced its outcome.

## Acknowledgements

PAH acknowledges Mike De Iuliis for continued support and encouragement. This work was supported by Macquarie University and the Biomolecular Discovery Research Centre.

## Notes

http://www.bitbucket.org/peptidewitch/peptidemind

### Competing Interest Statement

The authors have declared no competing interest.

